# Spatiotemporal dynamics between interictal epileptiform discharges and ripples during associative memory processing

**DOI:** 10.1101/2020.07.22.216416

**Authors:** Simon Henin, Anita Shankar, Helen Borges, Adeen Flinker, Werner Doyle, Daniel Friedman, Orrin Devinsky, György Buzsáki, Anli Liu

## Abstract

We describe the spatiotemporal course of cortical high-gamma activity (HGA), hippocampal ripple activity and interictal epileptiform discharges (IEDs) during an associative memory task in 15 epilepsy patients undergoing invasive electroencephalography. Successful encoding trials manifested significantly greater HGA in hippocampus and frontal regions. Successful cued recall trials manifested sustained HGA in hippocampus compared to failed responses. Hippocampal ripple rates were greater during successful encoding and retrieval trials. IEDs during encoding were associated with 15% decreased odds of remembering in hippocampus (95% CI 6-23%). Hippocampal IEDs during retrieval predicted 25% decreased odds of remembering (15-33%). Odds of remembering were reduced by 25-52% if IEDs occurred during the 500-2000 ms window of encoding or by 41% during retrieval. During encoding and retrieval, hippocampal IEDs were followed by a transient decrease in ripple rate. We hypothesize that IEDs impair associative memory in a regionally and temporally specific manner by decreasing physiologic hippocampal ripples necessary for effective encoding and recall. Because dynamic memory impairment arises from pathological IED events competing with physiological ripples, IEDs represent a promising therapeutic target for memory remediation in patients with epilepsy.

**Summary:** Hippocampal interictal epileptiform discharges in hippocampus acutely impair declarative memory, potentially by hijacking physiological processes essential for encoding and recall.

## INTRODUCTION

Memory dysfunction affects 40–50% of epilepsy patients. In a community-based survey, patients ranked memory problems as the most important cognitive comorbidity of epilepsy, adversely affecting daily functioning, work participation, and school performance (Fisher *et al*., 2000). These subjective complaints range from difficulties with remembering names and phone numbers (semantic memory) to what happened during last year’s family vacation (episodic memory) (Lemesle *et al*., 2017). While multiple factors are involved (Mazarati, 2008), interictal epileptiform discharges (IEDs) are pathological bursts of neuronal activity between seizures which can dynamically impair cognition. In 1939, Schwab demonstrated that ‘subclinical EEG discharges’ increased reaction time or resulted in failure to respond to the stimulus (Schwab, 1939). Subsequently, Aarts introduced the term ‘transitory cognitive impairment’ (TCI) to describe this functional disruption. Because IEDs are transiently associated with memory impairment, they may provide a target for therapeutic intervention (Aarts *et al*., 1984).

Intracranial EEG (iEEG) studies with epilepsy surgery patients have advanced our understanding of human cognition, including the deleterious impact of IEDs. iEEG offers high spatiotemporal precision and superior signal-to-noise ratio (Lachaux *et al*., 2012) for characterizing the neurophysiology of memory, including recordings from deep structures (e.g., hippocampus and entorhinal cortex) inaccessible by other methods. Increased gamma activity in dominant mesial temporal and frontal regions predicts successful encoding of word lists (Sederberg *et al*., 2003; Sederberg *et al*., 2007) or word pairs (Greenberg *et al*., 2015). Recently, hippocampal sharp-wave ripple events have been correlated with successful retrieval of word pairs (Vaz *et al*., 2019; Vaz *et al*., 2020) and visual episodic memories (Norman *et al*., 2019). However, these iEEG studies describing physiological processes including gamma activity and ripples typically exclude trials with IEDs, discarding them as “noise.”

Accumulating evidence demonstrates that IEDs can impair encoding, maintenance, consolidation, and retrieval of verbal learning (Aarts *et al*., 1984; Krauss *et al*., 1997; Stemmer *et al*., 2013; Gelinas *et al*., 2016; Horak *et al*., 2016; Ung *et al*., 2017). Left temporal and parietal neocortical IEDs are associated with impaired memory for word list items and word pairs (Horak *et al*., 2016; Ung *et al*., 2017). IEDs outside the seizure onset zone (SOZ) in higher order visual processing regions have been associated with impaired encoding and retrieval performance for words (Ung *et al*., 2017). Despite this, our understanding of the relationship of IEDs and human memory is largely correlational (Stemmer *et al*., 2013; Horak *et al*., 2016; Kleen and Kirsch, 2017), without a clear mechanism of disruption. Studies assessing word-list learning or verbal paired associate tasks found the greatest effects of IEDs occurring in higher-order visual neocortical areas (i.e. fusiform gyrus, inferior temporal gyrus) or parietal lobe (Horak *et al*., 2016; Ung *et al*., 2017) It is unclear whether IEDs disrupt sensory processing of words or memory function per se and if they do, the mechanism of disruption is unknown (Kleen and Kirsch, 2017). IEDs may be more frequent during drowsy or distracted states, and thus only indirectly associated with poor memory performance (Leung, 1988; Kleen and Kirsch, 2017).

In this study, we aimed to establish the relationship between high gamma activity and hippocampal ripples and pathological interictal epileptiform discharges (IEDs). Previously, physiological and pathological events have been investigated independently. We hypothesized that characterizing the spatiotemporal course of physiological activity during a memory task would reveal when and where in the brain IEDs exert the greatest impact and provide a potential mechanism for how IEDs disrupt cognitive processing. We selected a face-profession association task, which potentially represents a more clinically relevant probe of episodic memory than word-list recall or word-pair association tasks (Sederberg *et al*., 2003; Sederberg *et al*., 2007; Stemmer *et al*., 2013; Greenberg *et al*., 2015; Jang *et al*., 2017; Ung *et al*., 2017). Our face-profession task depends on bilateral hippocampal (Staresina and Davachi, 2006; Theysohn *et al*., 2013; Suthana *et al*., 2015) function, thus permitting an opportunity to test how IEDs, often abundant in the mesial temporal lobe, interrupt mnemonic processes. We examined the spatiotemporal time course of high gamma activity (HGA), a robust index of local cortical activity (Mukamel *et al*., 2005) across widespread brain areas during encoding and retrieval, distinguishing between successfully remembered versus forgotten pairs. As hippocampal activation was critical for encoding and retrieval, we next compared the hippocampal ripple rate of successful versus failed trials. To ensure that detected HGA and ripple rate were physiological events and not pathological high frequency oscillations, which are increased in epileptogenic cortex (Weiss *et al*., 2013; Weiss *et al*., 2016) and often coupled with IEDs (Weiss *et al*., 2016), we excluded electrodes inside the seizure onset zone (SOZ) and trials with IEDs. Next, we evaluated the impact of IEDs by brain region and with respect to the seizure onset zone (SOZ). Because hippocampal IEDs had a consistently adverse effect on memory, we examined their impact by time course, predicting a greater impact on memory if IEDs occurred during the trial when hippocampal activity was critical. Finally, to investigate how IEDs may disrupt memory function, we examined the temporal relationship between IEDs and hippocampal ripples.

## MATERIALS AND METHODS

### Face-profession association task

We employed 120 color images of distinct human faces with neutral expression from the Chicago Face Database (Fig. 2A)(Ma *et al*., 2015). The image set comprised 59 male, 61 female, and an equal proportion of White, Black, Hispanic, and Asian faces, paired with 120 emotionally neutral, single-word professions, 4–10 letters long, selected from the US Bureau for Labor Statistics database. Since epilepsy patients typically have impaired performance compared to healthy subjects, (Krauss *et al*., 1997; Fell *et al*., 2002; Stemmer *et al*., 2013; Horak *et al*., 2016) we calibrated the task difficulty per subject to achieve a balanced distribution between successful and failed encoding trials. Trial sets ranged from 1 to 10 pairs per set. Task stimuli were presented by computer using custom software (MATLAB, Psychophysics Toolbox). Each face-profession pair was shown for 5 seconds, with a 1 second interstimulus interval which was marked by a plus sign. To ensure attention and sensory processing of test stimuli, patients were instructed to read the profession aloud and make a mental association. To prevent rehearsal after the encoding block, a brief distraction task was presented (“Count backwards from 15”). The task laptop sent a pulse to a trigger input channel in the EEG amplifier to synchronize task stimuli with the electrophysiological recordings. The time stamps associated with the pulses were used to annotate the iEEG recordings. For the last 10 patients, we recorded audio responses, enabling analysis of HGA and hippocampal ripples aligned temporally with the onset of the patient’s vocalized response.

We tested for memory function using a cued recall paradigm. This paradigm allows greater experimental control compared to free recall designs, including the ability to precisely measure the spatiotemporal relationship between physiological and pathological events. A cued recall paradigm may be more sensitive to clinical memory dysfunction compared to recognition memory tasks (Cassel *et al*., 2016). During recall, subjects were shown only faces from the prior set and asked to speak aloud the associated profession. The cued recall segment lasted for as long as the subject needed to provide a response. Sets during which subjects freely stated the associated profession paired with the face stimulus were scored as correct trials. Trials during which the subject responded “pass” or gave the incorrect answer were scored as incorrect trials.

Antiseizure medications were reduced to record seizures during the patient’s hospital stay. Cognitive testing was performed >= 6 hours after the last seizure. If a patient had a seizure or non-epileptic seizure during testing, all concurrent and subsequent trials were excluded from analysis. All participants provided informed consent with procedures approved by the institutional review board at NYU Langone.

### iEEG data acquisition and electrode localization

Brain activity was recorded from implanted stainless steel or platinum-iridium depth electrodes or subdural electrode grids embedded in silastic sheets. Decisions for implantation, placement of electrodes, and the duration of monitoring were made by the clinical team and without reference to this study. Subdural grids and strips covered extensive portions of lateral and medial frontal, parietal, occipital, and temporal cortices of both hemispheres (Fig. 1). Recordings were made using: (i) Natus NicoletOne C64 clinical amplifier, bandpass filtered from 0.16–250 Hz, with a 512 Hz sampling rate or a (ii) Natus Quantum clinical amplifier, with a 2000 Hz sampling rate, which was later down sampled to 512 Hz following anti-aliasing filtering. Electrode localization was performed using automated processes and expert review. The location of each electrode relative to the cortical surface was determined from post-implantation CT scan or MRI Brain co-registered to the pre-implant T1-weight MRI Brain scan (Yang *et al*., 2012). Co-registered, skull-stripped T1 images were nonlinearly registered to an MNI-152 template and electrode locations were then extracted in Montreal Neurological Institute (MNI) space. Each electrode was assigned to a brain Region of Interest (ROI) based on the Desikan-Killiany anatomical atlas (Desikan *et al*., 2006). Hippocampal depth electrode location was confirmed by expert review (AL, SH).

**Fig. 1.**
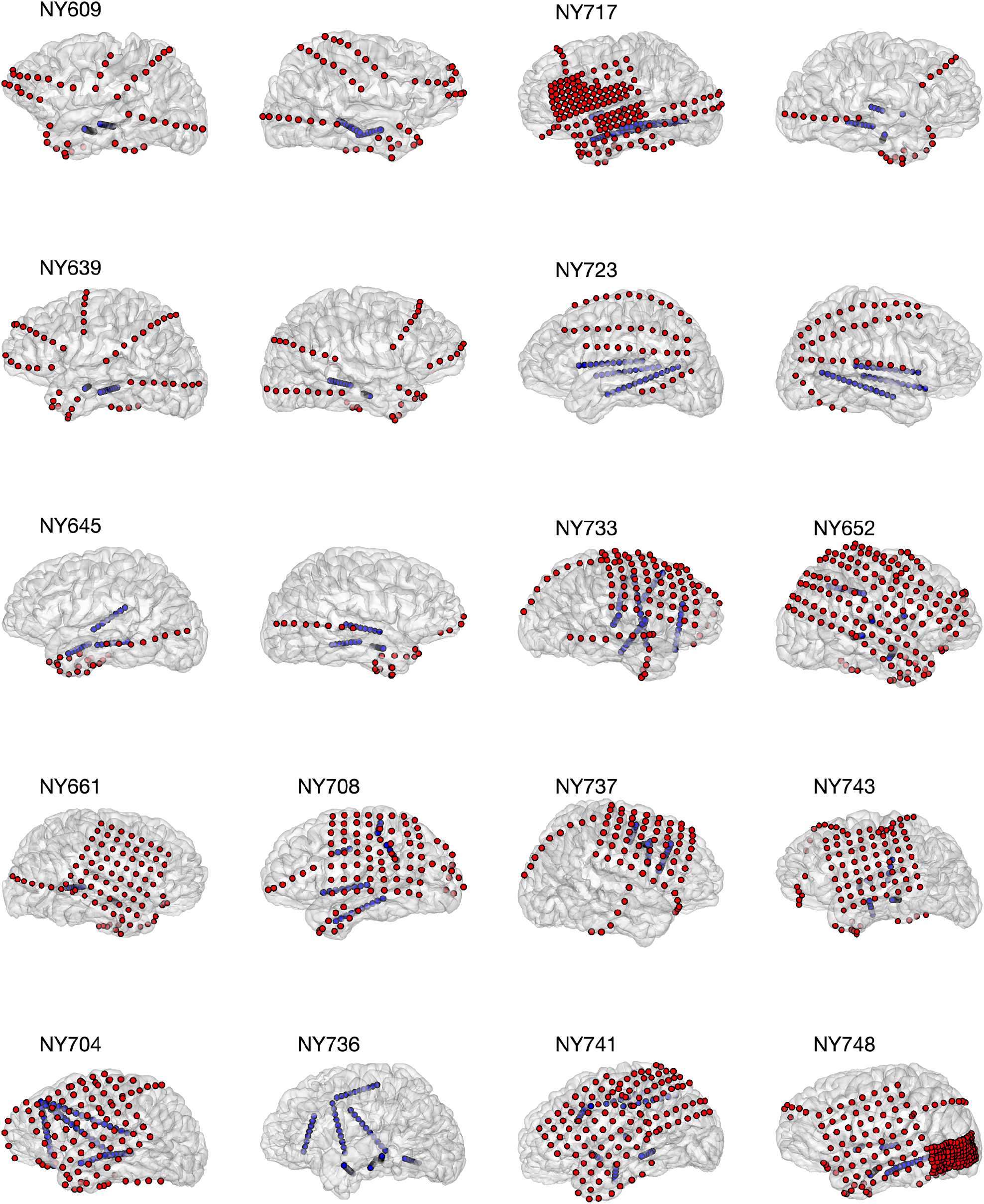
Electrode coverage for Fifteen Subjects. We recorded from a total of 1646 electrodes in 15 subjects. Five patients had bilateral depths and strips (NY609, NY639, NY645, NY723, NY736). Six patients had left hemisphere subdural grids, strips, and depths (NY704, NY 708, NY717, NY741, NY743, NY748). Four patients had right hemisphere subdural grids, strips, and depths (NY652, NY661, NY733, NY737). Grid and strip electrodes are shown in red; depth electrodes are shown in blue. NY737 did not have hippocampal depth electrodes and therefore was excluded from ripple analysis.

### iEEG analysis

Standard artifact rejection techniques, such as notch filtering for line noise and harmonics (60, 120, 180 Hz), detrending, and baseline correction were applied. In addition, EEG artifacts were removed by omitting trials where the raw signal exceeded 5 SD above the mean(Fell *et al*., 2001). Finally, visual inspection of individual trials excluded trials with excessive non-physiological noise. iEEG analysis focused on identifying differences in the spectral-temporal features between successful and failed encoding trials. All analyses were performed using a bipolar montage, conducted across regions and trials. After identifying electrodes in the hippocampus, entorhinal cortex, and regions of the neocortex, we performed time-frequency analyses in the −500 ms to 2000 ms from time of the cue presentation. For 10 subjects with recordings of spoken responses, we manually marked the onset of the vocalization and derived vocal-aligned responses for each trial in −2000 ms to 2000 ms around the vocal onset.

iEEG recordings were segmented into encoding and cued recall periods, comparing trials which were later successfully recalled versus forgotten. For recall, we performed two analyses: one aligned to cue presentation and another aligned against vocalization. HGA activity of the raw EEG traces was estimated using multi-taper spectral analysis (Chronux Toolbox, 250 ms windows, 10 ms steps; 60-170 Hz), normalized per frequency bin to the −0.5 to −0.05 s pre-stimulus baseline, and then averaged across frequency bands. For vocalization-aligned responses, baseline normalization was performed across the entire −2 to +2s interval from vocalization onset to adjust for the large variability in response time relative to cue onset. For each patient, HGA across all electrodes was averaged within a region of interest (ROI) based on the Desikan-Killiany anatomical atlas (Desikan *et al*., 2006). Finally, to assess for differences between conditions (successful vs. failed encoding) within a given ROI, a non-parametric cluster-based permutation test was used to determine significance of any power changes between conditions and control for multiple comparisons (Maris and Oostenveld, 2007). Briefly, temporal clusters are identified in the test set (e.g. adjacent timepoints which exceed the critical t-value using a dependent samples t-test, two-sided). Next, a null distribution was formed from 1000 random permutations of the condition labels, and clusters were identified in this set using the same criteria. Clusters in the test set were deemed significant if the summed t-statistic in the observed cluster exceeded that of the maximum cluster in the null distribution 95% of the time (p<0.05, cluster-corrected).

### Seizure onset zone

We sought to identify electrodes within the seizure onset zone (SOZ), which has impaired function compared to non-SOZ regions (Ung *et al*., 2017). The SOZ was determined by the clinical team, by correlating clinical seizures with intracranial EEG seizures. SOZ electrodes were excluded from all analyses correlating hippocampal ripples and IEDs with memory performance, and when examining the temporal relationship between IEDs and ripples. We performed a secondary analysis of ripple rate in SOZ to assess its functional reserve.

### IED Detection

Given the significant inter-rater variability in IED annotation among experts (Jing *et al*., 2019), we used an automated IED detection algorithm (Janca *et al*., 2015) combined with validation by 2 experts (AL, SH). This algorithm identifies brief outliers in the signal envelope (Hilbert envelope of the bandpass filtered signal between 10-60 Hz) by adaptively modeling the distribution of the background activity (log-normal distribution of the signal envelope in 5s windows, 4s overlap), and determining if the signal voltage exceeds 3.3x the mode+median of the envelope of the modeled background activity (Janca *et al*., 2015). All spikes in the 0 ms to +2000 ms window were identified for each electrode for both encoding and recall trials, and compared between correct and incorrect trials.

### IED Analysis and Modeling

To assess the impact of IEDs on memory, we fit a mixed-effects model to the IED counts per region of interest (ROI). A generalized linear mixed-effects model (GLMM) with a logit link function permitted modeling of a binary outcome of each trial (remembered versus forgotten)(Ung *et al*., 2017), as, *logit*(*π*_*ij*_) *= β*_*0*_ + *β*_*1*_*IED*_*ij*_ + *b*_*i*_ where, *π*_*i*_ is the probability of successful recall of subject j in trial i, *IED*_*ij*_ is the number of spikes in trial j for subject i, and *b*_*i*_ ∼ *N*(*0, σ*^*2*^) is the random-effects intercept for each subject i, accounting for subject-specific variation in recall performance. Effect sizes (odds ratio, OR) and confidence intervals (CI) were calculated from logistic regression estimates of the fixed-effects (e.g. *OR = exp*(*β*_*1*_)). Significance of the model coefficients were determined using an F-test (Korn and Graubard, 1990) and FDR corrected across all ROIs. IEDs in more than one region were considered independently (Ung *et al*., 2017).

### Ripple Detection

All electrode pairs (bipolar montage) with at least one contact located within the hippocampus (including anterior and posterior regions) were selected for analysis for each patient. To reduce the likelihood of detecting pathological ripples, electrodes within the SOZ and trials with IEDs were excluded from initial ripple analysis. Ripple detection was performed by filtering the raw iEEG between 80-120 Hz, then extracting the power envelope using the Hilbert transform. Events where the envelope of the filtered response exceeded 2 SDs and measured 20-200 ms in duration were marked as ripple events(Vaz *et al*., 2019). These filter settings reduced the potential of detecting pathological high frequency oscillations (HFOs), which have a centroid peak between 150-200 Hz (Liu and Parvizi, 2019). HFOs are potential biomarkers of epileptogenicity and have been negatively correlated with HGA and memory performance (Liu and Parvizi, 2019). Ripple rate was calculated in 125 ms bins, as the number of ripple events detected per second. Ripple duration and spectral qualities were inspected. To examine differences in ripple rate between successful and failed encoding, the average ripple rate was computed across all electrodes for each patient, and a non-parametric cluster-based permutation test was used to identify the effect of condition (successful versus failed) across patients.

A secondary analysis examined the impact of IEDs on ripples during encoding, cued retrieval, and voice-aligned retrieval, by comparing the hippocampal ripple rates before and after the IED detection. First, all IEDs across all hippocampal electrodes were pooled and binned in 500 ms windows within encoding, recall, and voice aligned trials, from 500 ms before to 2000 ms after cue onset. Ripple rate was calculated in the 500 ms window before and after each IED was detected, excluding ripples within 50 ms before and after the IED to avoid detection of pathological HFOs. Ripple rates before and after IEDs were compared using a paired Wilcoxon’s signed rank test, then adjusted for multiple comparisons using false discovery rate (FDR) corrections (Benjamini and Hochberg, 1995). This measure tested the null hypothesis that the ripple rate before and after the IED were the same.

### Data Availability

The data that support these findings are available upon reasonable request from the corresponding authors.

## RESULTS

### Data Collection

We recorded from fifteen (15) patients with epilepsy undergoing iEEG monitoring for surgical evaluation from New York University Langone Hospital (NYULH). All study activities were approved by the NYU Langone Institutional Review Board and all patients provided informed consent to participate in the study. We included a total of 7 males and 8 females, with an average age of 30.5 (range 15-55, SD 12.4). Fourteen were right-handed and one was ambidextrous; average IQ was 98.8 (range 82-132, SD 15.6). Table 1 summarizes additional demographic and clinical characteristics. Most patients scored below normative values on a standardized verbal memory task, the Rey Auditory Verbal Learning Test (RAVLT, z=-1.5, SD=2.4, Table S1), and visual memory task, the Rey Ostereith Copy Figure Test (ROCFT, z= −2.5, SD=2.4, Table S1). Patients’ implantations included a combination of subdural grids, strips and depth (e.g. hippocampal) electrodes (Fig. 1). For one patient (NY737), electrode coverage did not include the hippocampus. We analyzed 1646 electrodes in this study. Seizure onset zones (SOZ) were diverse (Table 1). Eight patients underwent surgical resection, six were recommended for neurostimulation (RNS, VNS, or DBS), and one patient did not receive a surgical intervention due to failure to record clinical seizures.

**Table 1.** Demographic and Clinical Characteristics of Subjects. Fifteen (15) subjects were recruited from a single epilepsy center. Subjects had an average age of 30.5 years (range 15 to 55, SD 12.4), 53% female, right-handed, with a mean IQ of 98.8 (Range 82-132, SD 15.6). Patients were implanted with a combination of strategies: bilateral strips and depths; right and left hemisphere grids, strips, and depths. Seizure onset zones determined by intracranial EEG monitoring and clinical outcomes are reported, which could include surgery or a therapeutic device (RNS: Responsive Neurostimulation, DBS: Deep Brain Stimulation or VNS: Vagal Nerve Stimulation).

|  | Subj | Age | Sex | Handness | Avg. Set Size | IQ | Coverage | Seizure Onset Zone (SOZ) | Outcome |
| --- | --- | --- | --- | --- | --- | --- | --- | --- | --- |
| 1 | NY609 | 45 | F | R | 2.2 | 95 | Bilateral | R mesial temporal | R anterior temporal lobe resection |
| 2 | NY639 | 18 | M | R | 8.8 | 91 | Bilateral | L hemisphere | RNS |
| 3 | NY645 | 38 | M | R | 4.3 | 82 | Bilateral | R mesial temporal | RNS |
| 4 | NY652 | 20 | F | R | 4.1 | 74 | R hemisphere | R temporal, parietal, occipital | R temporal, parietal, occipital corticectomy |
| 5 | NY661 | 28 | M | R | 9.2 | 132 | R hemisphere | R temporal | R anterior temporal lobe resection |
| 6 | NY704 | 55 | F | R | 3.5 | 108 | L hemisphere | L basal and lateral temporal neocortex | Tailored L anterior temporal lobe resection |
| 7 | NY708 | 41 | F | R | 1.4 | 116 | L hemisphere | Unclear<br>Non-epileptic seizures | No surgical intervention |
| 8 | NY717 | 49 | F | R | 3.3 | 115 | Bilateral | R temporal pole and<br>hippocampus | RNS |
| 9 | NY723 | 26 | M | R | 2.4 | NA | Bilateral | B mesial temporal lobes | RNS |
| 10 | NY733 | 15 | M | R | 4 | 107 | R hemisphere | R orbitofrontal, cingulate,<br>insula | Tailored R frontal<br>corticectomy |
| 11 | NY736 | 29 | M | R | 4 | 96 | LH depths | L mesial temporal cortex | L anterior temporal lobe<br>resection |
| 12 | NY737 | 20 | F | R | 4 | 91 | L hemisphere | R middle frontal, cingulate,<br>pars triangularis/opercularis,<br>precentral | R frontal corticectomy |
| 13 | NY741 | 24 | M | B | 2.1 | 82 | L hemisphere | L frontal, parietal, temporal,<br>occipital | RNS, VNS, or DBS |
| 14 | NY743 | 30 | F | R | 2 | 98 | L hemisphere | L posterior insula,<br>periopercular | RNS |
| 15 | NY748 | 19 | F | R | 3.4 | 96 | L hemisphere | L temporal neocortical | Tailored L post inferior<br>temporal corticectomy |

All patients performed the face-profession visual association task (Fig. 2A). The face-profession task robustly activates both hippocampi in healthy subjects (Theysohn *et al*., 2013). Since epilepsy patients typically have impaired performance compared to healthy subjects, (Krauss *et al*., 1997; Fell *et al*., 2002; Stemmer *et al*., 2013; Horak *et al*., 2016) we calibrated the task difficulty for each subject to achieve a balanced distribution between successful and failed encoding trials. Subject performance varied from one to ten trials per set, between 24 to 119 trials total per subject. Low total trial numbers were due to poor performance, lethargy, ensuing seizure, and/or time limitations due to clinical factors. Because trial set size varied by subject, direct comparisons of performance across subjects was not possible.

**Fig. 2.**
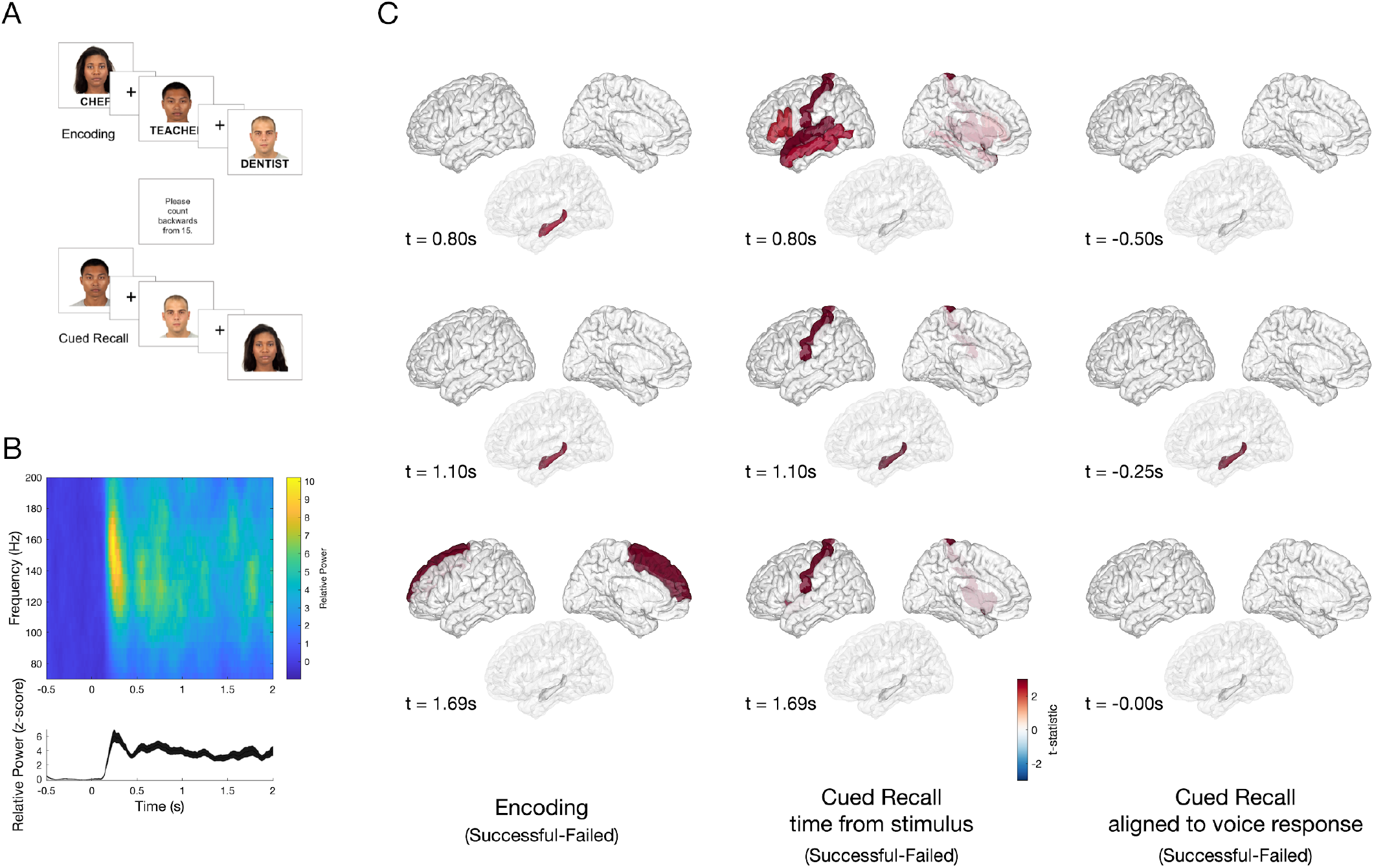
Task Design and Differences in High Gamma Activity (HGA) between Successful and Failed Associative Memory. **(A)** A computerized program presented task stimuli and recorded subject spoken responses. During encoding, each face-profession pair was shown for 5 seconds, with a 1 second inter-stimulus interval (ISI) which was marked by a plus sign. To ensure attention and sensory processing of test stimuli, subjects were instructed to read the profession aloud and make a mental association. To prevent rehearsal, a brief distraction task followed the encoding block, during which subjects were asked to count backwards from 15. During cued recall, subjects were shown only the faces from the prior set and asked to say aloud the associated profession. The cued recall period lasted for as long as the subject needed to provide a response. Voice response was recorded for the last 10 subjects and scored for accuracy. **(B)** Example spectrogram (top) of the raw data recorded in the occipital cortex, and high gamma activity (bottom, HGA 60-170 Hz) normalized to the −500 pre-stimulus baseline, with a peak at 250 ms after stimulus presentation. (**C**) Group-level differences in HGA by time for correctly versus incorrectly recalled face-profession pairs thresholded at p<0.05 (cluster-corrected) during encoding (left), cued recall (middle) and vocal-aligned cued recall (right). **Left**. During encoding, increased HGA in hippocampus beginning approximately +0.80s after stimulus presentation, with increased HGA in superior frontal region beginning approximately +1.69 s distinguished between successful and failed trials. **Middle**. During cued recall, increased HGA at +0.80s after face stimulus presentation in inferior frontal gyrus, postcentral, superior temporal and middle temporal gyrus, and later at +1.10s in hippocampus distinguish between successful and failed trials p<0.05, cluster-corrected). **Right**. To disambiguate the contribution of vocalization to cued recall, the difference between successful and failed trials was determined, timed in response to the vocalization in 10 patients. A difference in hippocampal HGA was seen beginning at −250 ms prior to vocalization (all significant clusters identified at a significance threshold p<0.05 using a cluster-based permutation test).

### Differences in High Gamma Activity in Hippocampus and Frontal Regions Distinguish Successful versus Failed Remembering

iEEG recordings were segmented into encoding and cued recall periods, comparing trials which were later successfully recalled versus forgotten. As expected, high gamma activity (HGA, 60-170 Hz) demonstrated an early increase in several cortical areas, including primary visual cortex (**Fig 2B**) and visual association cortices, such as fusiform gyrus and inferior temporal gyrus, immediately after stimulus presentation (**Fig. 2C**). Later, beginning approximately + 0.80s after stimulus presentation, HGA in hippocampus distinguished successful and failed encoding trials (p<0.05, cluster-correct, additionally, see Fig. S4). Increased HGA in the superior frontal region beginning at +1.69 s also characterized successful encoding (Fig. 2C, p<0.05, cluster-corrected). During cued recall, increased HGA at +0.80s after stimulus presentation in inferior frontal, postcentral, superior and middle temporal gyri, and later at +1.10s in hippocampus and postcentral gyrus distinguished successful and failed recall trials (Fig. 2C, p<0.05, cluster-corrected). To disambiguate the cognitive from the motor vocalization response components and to account for variable response times, we examined the difference between correct and incorrect responses timed to the vocalization in 10 patients with voice recordings. We found a difference in hippocampal HGA occurring before vocalization, starting approximately −250 ms before vocalization (Fig. 3C, p<0.05, cluster-corrected).

**Fig. 3.**
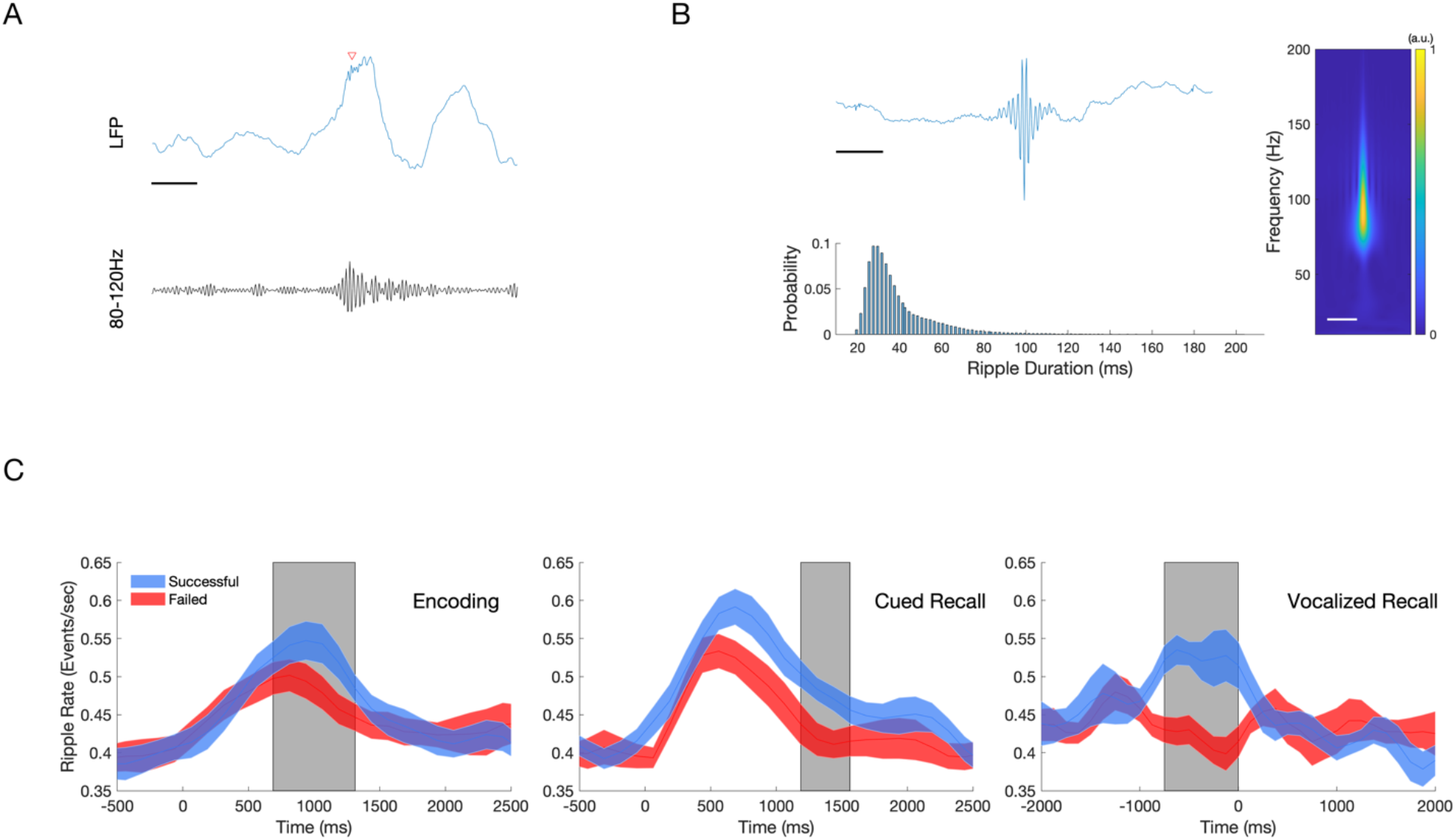
Hippocampal Ripples During Encoding and Recall Predict Successful Associative Memory. Ripple events were detected using a bipolar montage from the electrodes located in or closest to the hippocampus, using a previously published method (80 Hz-120 Hz, 20-200 ms duration, (Vaz *et al*., 2019)). To reduce the detections of pathological high frequency oscillations (HFOs), detections were restricted to regions outside of seizure onset zone, and with trials which did not contain an interictal epileptiform discharge. **(A)** Sample raw EEG tracing (blue) with detected ripple event (RED ARROW), with bandpass filtered (80-120 Hz) tracing (black). Scale bar, 125ms **(B)** Characteristics of all detected hippocampal ripples. An average of 1955 ripple events were detected per patient across all conditions. **Top:** Grand averaged ripple response (left) and spectrogram (right, 10-200Hz) demonstrates a peak frequency between 80-100 Hz. Scale bars, 125ms. **Bottom:** Histogram showing detected ripple duration, which follows skewed log distribution. Mean ripple duration 39.8 ms, SD 18.3 ms. **(C)** Average ripple rate between Successful and Failed Associative Memory Trials (mean +/-SEM). **Top:** Successful encoding is characterized by a higher hippocampal ripple rate (blue) compared to failed encoding (red) between 750-1375ms after stimulus presentation (gray box, N=14, P<0.05, cluster-corrected). **Middle:** Successful cued recall is characterized by a higher ripple rate compared to failed cued recall between +1250 ms and +1625 ms after stimulus presentation (gray box, N=14, P<0.05, cluster-corrected). **Bottom:** Successful cued recall is characterized by a higher ripple rate from −750 ms to 0 ms aligned voice response (gray box, N=9, P<0.05, cluster-corrected).

### Hippocampal Ripples Increased in Successful versus Failed Remembering

Hippocampal ripples were measured from all depth electrodes within the hippocampus for each patient. Ripple detection was performed using a published method that has demonstrated the relationship between ripples in hippocampus and middle temporal gyrus and successful retrieval (80-120 Hz, 20-200 ms duration)(Vaz *et al*., 2019). We detected an average of 1955 (±799) hippocampal ripple events across all test blocks per patient, after excluding electrodes in the SOZ. An example raw trace of a detected ripple event is shown in Fig. 3A. We observed that ripples are brief events possessing oscillatory features characterized by a spectral centroid occurring between 80-120 Hz (Fig. 3B). Ripples had a median duration of 39 ms (±18 ms, Interquartile range) and showed a skewed log distribution (Fig 3B, bottom panel).

To examine whether the incidence of detected hippocampal ripple events during encoding and retrieval stages correlated with memory performance, we computed the ripple rate (Hz) of detected events in 125-ms bins from −500 ms to 2000 ms relative to the picture onset. While baseline ripple rates in the 500 ms preceding stimulus presentation were similar preceding all trials, successful encoding trials were characterized by a greater ripple rate approximately +750 to +1375 ms after stimulus presentation (Fig. 3C, top, p<0.05, cluster-corrected). Likewise, successful recall trials were characterized by a greater increase in ripple rate occurring approximately +1250 until +1625 ms after stimulus presentation (Fig. 3C, middle, p<0.05, cluster-corrected). For the ten patients with voice recording, successful trials had significantly increased ripple events in the 750 ms window preceding the vocal response (Fig. 3C, bottom, p<0.05, cluster-corrected).

In contrast, hippocampal ripples in the SOZ during encoding and cued recall did not differ between successful and failed trials (Fig. S2). However, even in the SOZ, ripple rate differentiated successful and failed trials when trials were aligned to the vocal response (Fig. S2).

### IEDs in Hippocampus Decrease Odds of Remembering

IEDs occurring during either the encoding or retrieval stages of the associative memory task were identified and pooled by anatomical region. We detected 13,484 IEDs across all test blocks and electrodes for all patients (mean and SD, 898 ± 342 events). IED rate varied from 0-2 events per trial across subjects (Fig. S5). We blindly reviewed approximately 12% of detected events, classifying them as a true or a false IED detection (AL, SH). This analysis revealed a positive predictive rate of 92% and false positive rate of 8% among the detected IEDs. **Fig. 4A** shows the raw tracing of an example IED and its spectrogram.

**Fig. 4.**
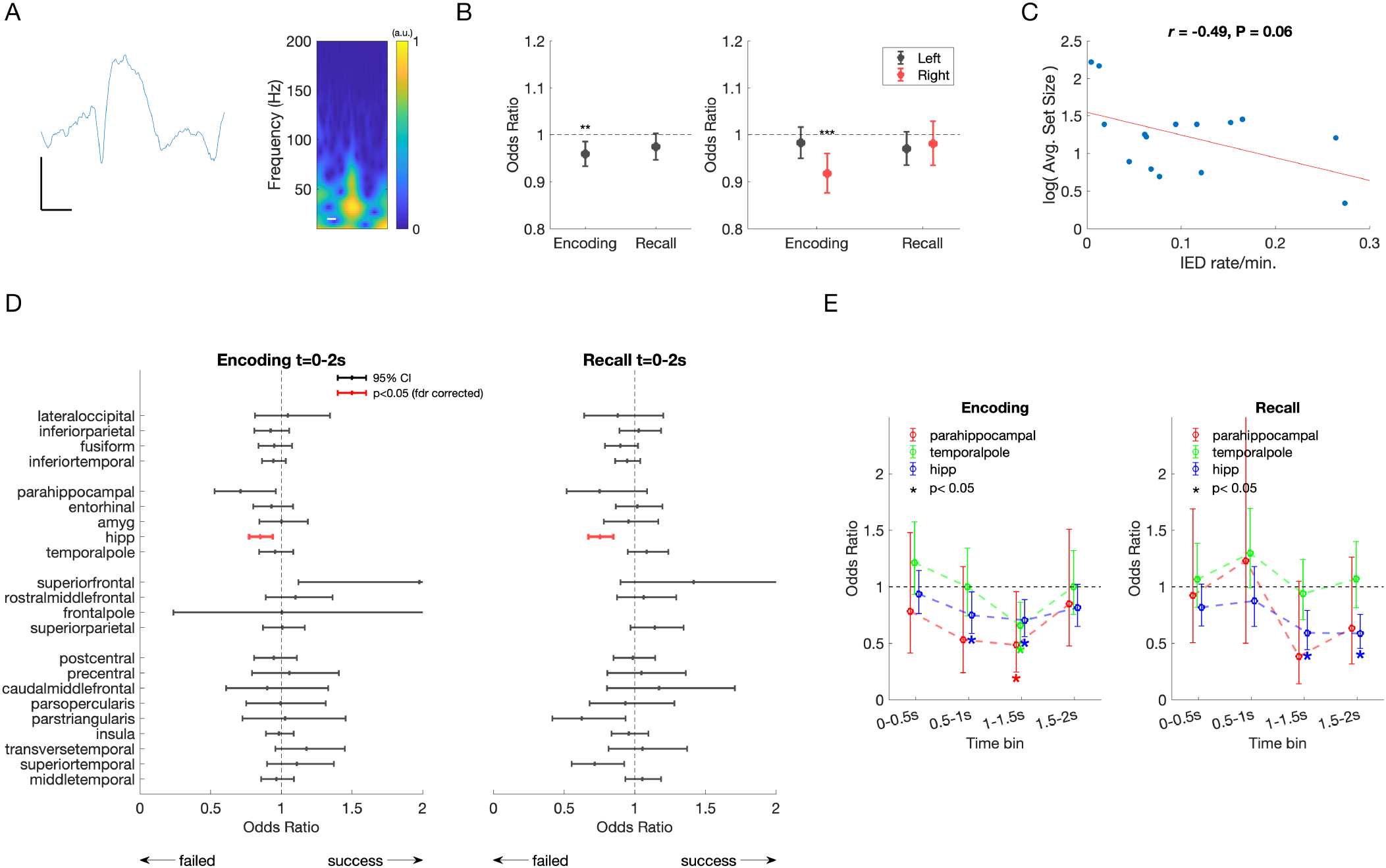
Interictal epileptiform discharges (IEDs) and effect on memory performance. **(A)** Example raw tracing and spectrogram of a detected IED. Scale bars, 25uV, 125ms **(B)** LEFT. IEDs recorded in any brain region during encoding predicted a 4% decreased odds of remembering (OR = 0.96, CI = (0.93-0.99), P = 0.002, F-test). IEDs in any brain region during cued recall trended toward decreased odds of remembering (P = 0.0760, F-test). RIGHT. IEDs occurring in the right hemisphere predicted a 9% decreased odds of remembering (P = 0.0002, F-test). There was no difference in odds of remembering for IEDs in the left hemisphere during encoding, or either hemisphere during recall. Error bars represent 95% confidence intervals **(C)** Relationship between block size, IQ, and IED rate. Subjects varied in performance, ranging between 1 to 10 stimuli presented per block, larger sets indicating superior task performance across patients. There was a trend toward a negative correlation between IED rate and log set size (*r*_13_ = −0.49, P = 0.06) **(D)** Odds of successful memory for IEDs occurring during encoding and recall, by brain region. Mean odds of successful memory per IED occurring during Encoding (LEFT) and Cued Recall (RIGHT). Error bars represent 95% confidence intervals. Odds less than 1 indicate a decreased odds of successful remembering if an IED occurred during the trial. After correction for multiple comparisons, a significant decrease in remembering occurs for IEDs in hippocampus (red, Encoding: OR = 0.85, CI = (0.77-0.94), F_1, 6221_ = 10.5, P = 0.001; Recall: OR = 0.75, CI = (0.67-0.85), F_1, 5956_ = 21.8, P < 0.001). **(E)** Odds of successful memory for IEDs occurring in selected brain regions, by 500 ms time bin. Mean odds of successful memory per IED occurring during Encoding (LEFT), demonstrate that odds of remembering are further decreased by 25-52% if IEDs occurred in hippocampus, parahippocampal gyrus, and temporal pole occur between 500-2000 ms. During recall (RIGHT), mean odds of successful memory per IED in hippocampus decreased by 41% when IEDs occurred between 1000-2000 ms after stimulus presentation. Error bars represent 95% confidence intervals. Significant changes indicated by an asterisk (P < 0.05, F-test).

SOZ was determined clinically by correlating seizure behavior to ictal iEEG changes. IEDs occurring in any brain region outside the SOZ during encoding decreased the odds of remembering (**Fig. 4B**, left, OR = 0.96, CI = (0.93-0.99), F_1,136290_ = 9.2247, p=0.002). IEDs occurring in any brain region during the cued recall stage was associated with a trend towards poorer performance (**Fig. 4B**, F_1,131202_ = 3.1477, p=0.08). Further, an IED occurring in the right, but not left, hemisphere during encoding was associated with significantly decreased odds of remembering (**Fig 4B**, right, OR = 0.92, CI = (0.88-0.96), F_1,61266_ = 13.8, p<0.001). There was a non-significant negative correlation between the frequency of IEDs and trial set size (r_13_=-0.49, p=0.06, Fig. 4C). Verbal IQ did not predict trial set size.

When evaluating the odds of remembering related to IEDs occurring in distinct brain regions during the encoding period, the largest impact on performance was when IEDs occurred in the hippocampus. An IED in the hippocampus was associated with decreased odds of remembering by 15% (95% CI: 6%-23%, Fig. 4D). During cued recall, an IED in hippocampus and outside of SOZ reduced odds of remembering by 25% (15-33%, Fig. 4D).

Given the time course of HGA and hippocampal ripple activity predicting successful versus failed trials, we predicted that IEDs during critical time windows would have an even greater effect. When the impact of IEDs during encoding was evaluated in a more time-resolved manner, we found that an IED in hippocampus occurring during the 500 ms to 1500 ms window after stimulus presentation was associated with decreased odds of remembering between 25% to 30% (Fig. 4E). IEDs in the temporal pole during the 1500 ms to 2000 ms window decreased odds of remembering by 34%. IEDs in parahippocampal cortex during the 1000-1500 ms after stimulus were associated with a decreased odds of 52%. Similarly, during cued recall, IEDs in hippocampus during the 1000-2000 ms window decreased odds of recall by 41% (Fig. 4E).

### Hippocampal IEDs Slow Response Time

In general, response times for all correct trial responses were delivered more quickly compared to incorrect responses or responses where subjects “passed” (mean = 1.74 s ±1.6 SD for correct trials vs. mean = 5.87 s ± 4.6 for incorrect/pass trials, Z=-14.2, p<0.001, Wilcoxon’s rank sum test) (**Fig. S6**). For all response types (correct, incorrect, pass), trials with an IED were slower compared to trials without an IED when trials were pooled across patients (**Correct:** median_IED-_ = 1.25 s (1.10 IQR); median_IED+_ = 1.61 s (1.49 IQR), Z = −3.89, p<0.001; **Incorrect:** median_IED-_ = 2.05 s (2.62 IQR); median_IED+_ = 4.15 s (3.93 IQR), Z = −5.19, p<0.001, **Pass:** median_IED-_ = 3.01 s (4.03 IQR); median_IED+_ = 6.13 s (4.35 IQR), Z=-3.74, p<0.001; Wilcoxon rank sum test).

### Hippocampal IEDs Decrease Ripple Rate

Because both IEDs and ripple rates were inversely related with memory performance, we examined their temporal relationship on a finer temporal scale. Hippocampal ripples generally decreased after the IED for during encoding and cued recall (Figs. 5A & 5B). However, this decrease in ripple rate reached significance during the 0.5-1.0 s and 1.0-1.5 s time bins during encoding. Additionally, ripple rate demonstrated the greatest decrease after IEDs occurring in the time bin aligned to vocal onset during cued recall (Fig. 5C). We further analyzed ripple rate decreases following IEDs along the longitudinal axis of the hippocampus, and found that IED induced reductions in ripple rates were primarily driven by activity in anterior hippocampus (Fig. S7).

**Fig. 5.**
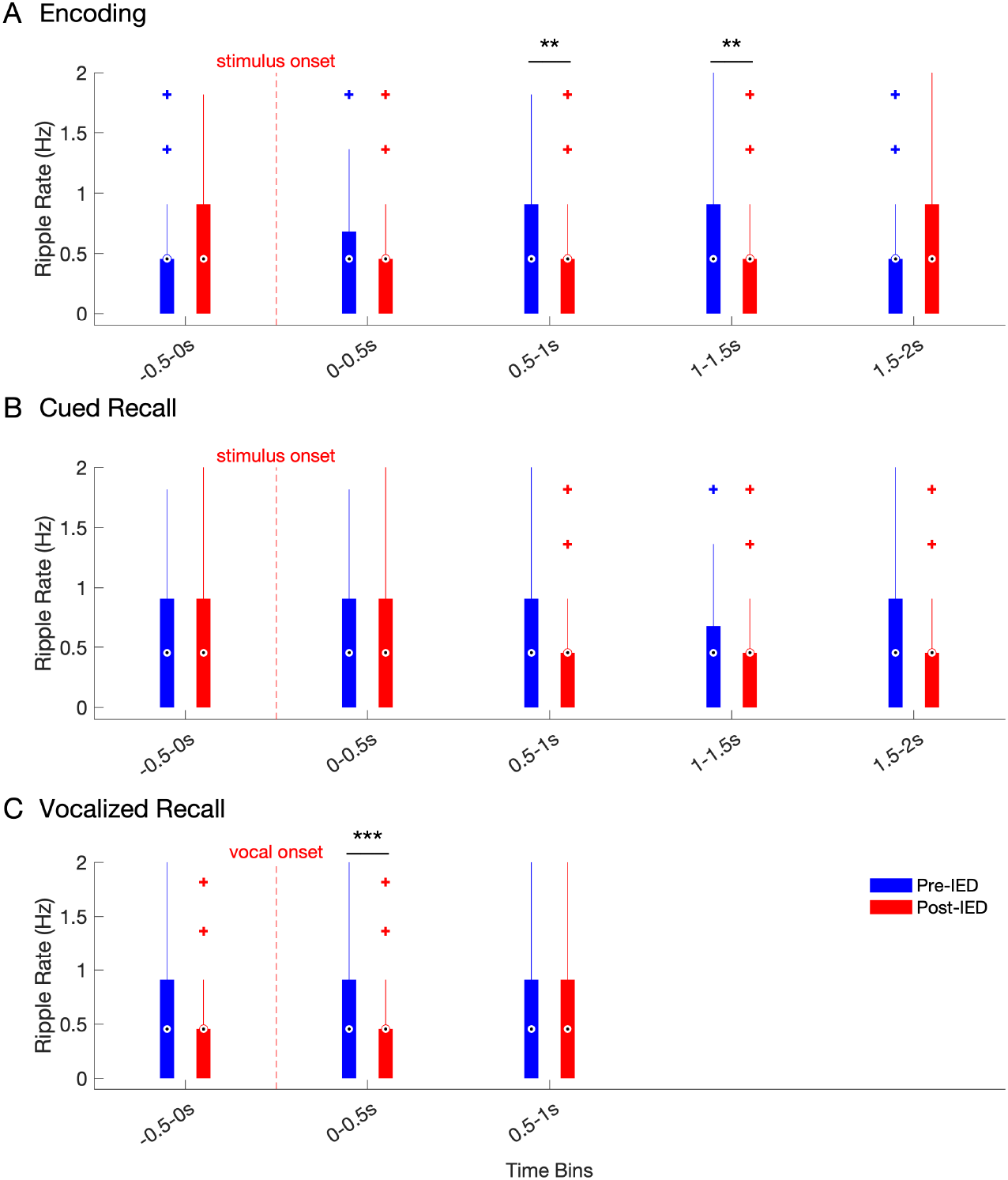
Hippocampal Ripple Rate in the Pre and Post-IED window during Encoding & Recall. Ripple rates across all hippocampal electrodes in 500ms pre-(blue) vs. 500ms post-IED (red), binned by time of detected IEDs (500ms bin windows) for IEDs detected during **(A)** Encoding, **(B)** Cued Recall, and **(C)** Voice-aligned Recall. Box and whisker plots represent the median (circles) and interquartile range (bars), along with extreme values (whiskers and outliers). A reduction in ripple rate after an IED event was found during the 0.5-1s (Z=2.8838, N_IEDs_=209, p=0.004) and 1-1.5s (Z=3.0873, N_IEDs_=266, p=0.002) windows during encoding. In addition, ripple rates were significantly reduced in the time window during (0-0.5s) vocalization during cued recall (Z=3.9, N_IEDs_=177, p<0.001).

## DISCUSSION

We examined the contribution of physiological (HGA and hippocampal ripples) and pathological (IED) events in successful and failed associative memory in epilepsy patients. An increase in frontal and hippocampal high gamma activity and hippocampal ripple rate during encoding predicted successful memory. During the encoding and cued recall stages, physiological differences between successful and error trials peaked between 500-1500ms after stimulus presentation, or within 750 ms of the spoken response. Conversely, pathological IEDs occurring in hippocampus during encoding and retrieval reduced the likelihood of remembering. A hippocampal IED during encoding decrease the odds of remembering by 15% (CI 6-23%), A hippocampal IED during cued recall decreased the odds of remembering by 25% (CI 15-33%). IEDs occurring during the 500-2000 ms window of encoding or retrieval had an even greater negative behavioral impact. Hippocampal IEDs observed during encoding and recall were generally followed by a decrease in ripple rate, although this difference only reached significance between 500-1500 ms during encoding and 0 to +500 ms around vocal onset. Overall, our findings suggest that IEDs impair associative memory in a regionally and temporally specific manner, likely by competing with physiological memory processes (i.e. hippocampal ripples) needed to encode and recall.

### Hippocampal Ripples Predict Associative Memory Performance

Our findings advance understanding of the contribution of physiological ripples to memory performance. Hippocampal ripple events detected from local field potential (LFP) recordings in humans undergoing invasive monitoring for epilepsy surgery have been shown to correlate with successful retrieval of word pairs (Vaz *et al*., 2019; Vaz *et al*., 2020) and visual episodic memories(Norman *et al*., 2019). Recently, LFP ripples in the middle temporal gyrus have been shown to organize sequences of single unit firing activity in humans in a phase-locked manner during successful memory retrieval(Vaz *et al*., 2020). The sequence of single unit firing carries item-specific information, with the temporal order of unit firing during encoding replayed during retrieval in remembered trials(Vaz *et al*., 2020). Similar to prior studies (Norman *et al*., 2019; Vaz *et al*., 2019; Vaz *et al*., 2020), we also observed an increase in ripple activity during the 750 ms prior to voiced retrieval predicting successful memory. Together, these studies represent a translational link to ripple activity observed in rodents during spatial navigation, which are replayed during wakefulness (Jadhav *et al*., 2012) and sleep (Wilson and McNaughton, 1994; Buzsaki, 2015).

Whereas previous human studies demonstrated ripples’ contribution to retrieval processes, we extend these findings to demonstrate the role of hippocampal ripple events during successful encoding and retrieval. To capture physiological, and not pathological, high frequency oscillations, we excluded SOZ electrodes. We also excluded trials with IEDs, which often have superimposed pathological HFO events overriding the peak of the waveform (Bragin *et al*., 1999; Waldman *et al*., 2018). Concordant with previous results, we found that ripples possessed an oscillatory waveform, a centroid peak between 80-100 Hz, a mean duration of ∼30 ms, and followed a skewed log distribution. These features resemble ripples in rodents (Bragin *et al*., 1999) suggesting that the events are not filtered high gamma events. The detected ripple rate at our threshold was approximately 1 event per second, comparable to previous human studies (Norman *et al*., 2019; Vaz *et al*., 2019). Most importantly, physiological ripple events had a strong temporal correlation with successful encoding and recall.

Our finding that ripple rate from hippocampus within SOZ distinguished between successful and failed trials only during the voice-aligned recall condition, but not during encoding or cued recall, suggests that SOZ preserves some physiological function, albeit less than tissue outside SOZ (Liu and Parvizi, 2019).

### Hippocampal IEDs decrease memory performance and prolong reaction time

Our work supports and extends prior work on the role of IEDs in memory dysfunction by detailing IED spatiotemporal dynamics, especially regarding the role of the hippocampus. Early studies using intracranial depth recordings in humans demonstrated that hippocampal IEDs during a Sternberg (letter) working memory experiment impaired performance if they occurred during the maintenance or recognition stage (Stemmer *et al*., 2013). One study including 80 surgical patients across multiple surgical centers demonstrated that IEDs during the encoding and recall epochs impaired verbal memory performance, with stronger effects seen for left hemisphere IEDs occurring in inferior temporal, medial temporal, and parietal regions (Horak *et al*., 2016). However, hippocampal IEDs predicted reduced memory only if they occurred during the recall phase, but not encoding phase(Horak *et al*., 2016). Another study in epilepsy surgical patients showed that IEDs occurring in the left fusiform gyrus, and middle and inferior temporal gyrus, and outside of SOZ, during encoding of a word list decreased odds of recall by 8-15% per IED event (Ung *et al*., 2017). The authors hypothesized that IEDs disrupted visual encoding or recognition of word forms. In prior work, it is surprising that hippocampus and nearby structures have been inconsistently implicated in IED-related memory disruption given their critical role in temporarily binding new information for later retrieval (Squire, 1992; Leranth *et al*., 1996; Eichenbaum, 2013). It is unclear from previous studies why IEDs would disrupt one stage of learning but not the other within the same task.

We show consistent and large effects of IEDs in hippocampus and adjacent neocortical regions during both encoding and recall, amplified during physiologically important windows of the trial. Our large effect sizes may be due to the appropriate selection of a face-profession association task, which differs from working memory paradigms and verbal free recall in several ways: 1) the task is associative in nature and demands linking semantic and visual information (a key function of the hippocampus) (Squire, 1992; Eichenbaum, 2013), and 2) it is a cued recall paradigm, allowing for precise temporal alignment between HGA, ripple events, and IEDs to stimulus presentation and vocal response. Here, we examined how the HGA and hippocampal ripple time courses could predict when and where IEDs might have maximal impact. We found that IEDs in mesial temporal structures occurring within 500-1500 ms—around the same time when HGA and ripple activity distinguished successful from failed memory--had a larger deleterious effect, decreasing odds of recall by 25-52% per IED. This is a truly large effect, implying that IEDs occurring within critical time windows will interfere with the learning process with a high probability.

Not only did IEDs increase forgetting, but slowed reaction times. Slower responses could reflect disruption in associative computation, but alternatively slower processing of the visual stimuli or execution of the motor command, depending on the IED location. Previously, focal and generalized IEDs have prolonged reaction times in a driving simulation task (Krestel *et al*., 2011; Nirkko *et al*., 2016), and increased crash rate when virtual obstacle is presented (Nirkko *et al*., 2016) IEDs prolonged reaction time in a variable manner, with generalized discharges affecting RT greater than focal discharges (Nirkko *et al*., 2016); and longer and higher voltage IEDs exerting a greater effect (Cohen *et al*., 2020).

### Hippocampal IEDs outside the SOZ Impair Memory Performance

Similar to previous reports (Ung *et al*., 2017), we found that IEDs within the SOZ generally did not impair memory performance. Only IEDs outside of SOZ diminished performance. This suggests that physiological processes, such as ripples, were already compromised in the epileptogenic cortex. We found a positive, consistent relationship with hippocampal ripple activity and memory performance when SOZ tissue was excluded from analysis. However, when hippocampal tissue within SOZ was analyzed separately, we only observed the increase in ripple rate predicting performance with voice aligned recall, but not during encoding or cued recall (**Fig. S1**). Our findings suggest some preserved function in SOZ compared to surrounding brain tissue outside of the SOZ (Liu and Parvizi, 2019)(Fig. S1). The diminished reserve in SOZ is likely due to greater neuronal loss and damage (Sutula *et al*., 2003).

### Hippocampal IEDs decrease ripple rate

By relating the temporal evolution of IEDs to hippocampal ripples, we demonstrate a potential mechanism for IED-induced disruption of mnemonic processing, at least in the anterior hippocampus. Previous studies focused on either physiological HGA or ripple activity (Norman *et al*., 2019; Vaz *et al*., 2019) or pathological IEDs (Stemmer *et al*., 2013; Horak *et al*., 2016; Ung *et al*., 2017). We have shown that IEDs occurring within the critical 1000-2000 ms period after cue presentation corresponds to a decrease in ripple activity required for successful recall. While IEDs have previously been demonstrated to interfere with sleep-dependent memory consolidation (Gelinas *et al*., 2016), our work suggests that IEDs also interfere with the physiological processes supporting memory encoding and recall. IEDs likely trigger a prolonged cortical downstate (Gelinas *et al*., 2016), as supported by a recent study demonstrating IED modulation of inhibitory interneurons in the medial temporal lobe (Reed *et al*., 2020). The state-dependent decrease in ripple activity after IEDs imply that IEDs are not merely an epiphenomenon of a drowsy or distracted state (Kleen and Kirsch, 2017), but rather trigger decreases in physiological ripples necessary for memory.

### Limitations

The limitations of our study include a moderate sample size of patients, which limit analysis of how clinical and demographic factors--such as gender, epilepsy duration and severity, and education--affect memory performance. In addition, given that intracranial recordings are composed of LFPs, we could not directly measure ripple content. We relied on the features of the filtered ripple event such as duration and frequency, their correlation with memory performance, differing time course from HGA to deduce that the signals were indeed physiological ripples. Strict exclusion of pathological cortex and trials and prior studies combining LFP with single unit recordings (Vaz *et al*., 2020) provide additional support. Future studies may utilize a combination of LFP with single unit recordings to reveal ripple content. Finally, studies in human subjects are limited by the inability to induce timed IEDs and prove their causative role in disrupting physiological models to impact memory.

## Conclusions

Earlier animal studies found that most hippocampal IEDs act as pathological ripples, recruiting a much larger population of the neuronal pool and in a narrower time window than the physiologically protracted events during ripples (Riekkinen *et al*., 1991). The competition between physiological ripples and pathological IED rate in a kindling model led to deterioration of performance on a cheeseboard maze task (Gelinas *et al*., 2016). Together, these findings and our work suggest that epileptic IEDs hijack physiological processes essential for effective encoding and recall of episodic memories.

Assuming that transient memory impairment arises from pathological IED events competing with physiological ripples, hippocampal IEDs represent a promising therapeutic target for memory remediation in patients with epilepsy and Alzheimer’s Disease (Lam *et al*., 2017). However, before IEDs can be considered to be a therapeutic target, assessment of performance in a real-time manner using a wider variety of clinically meaningful memory tasks is needed. Furthermore, closed-loop termination of IEDs during cognitive testing in rodents and humans (Krook-Magnuson and Soltesz, 2015) are needed to prove that blocking IEDs can indeed restore memory.

## ACKNOWLEDGEMENTS

The authors would like to acknowledge Lucia Melloni and Lila Davachi for early advice on task design.

