## Supplemental Materials for "Spatiotemporal dynamics between interictal epileptiform discharges and ripples during associative memory processing"

**SUPPLEMENTARY TABLE AND FIGURES**

| Subject | RAVLT | | | | R-OCFT | | | | TOPF |
|---|---|---|---|---|---|---|---|---|---|
| | | | | | Copy | | Delay | | |
| | Raw | Z-Score | Hits | Z-Score | Raw | Z-Score | Raw | Z-Score | |
| NY609 | 22 | -3.4 | 10 | -2.9 | 29 | -2.2 | 0 | -3.7 | 107 |
| NY639 | | | | | 25 | | 20 | | |
| NY645 | | | | | | | | | 95 |
| NY652 | 34 | -3 | 14 | -0.2 | 31 | -2.3 | 16.5 | -1.8 | 87 |
| NY661 | 53 | 0.4 | 15 | 0.6 | 33 | -2.7 | 17.5 | -1.7 | 123 |
| NY704 | 44 | -0.4 | 15 | 0.8 | 28 | -4.5 | 12.5 | -1.1 | |
| NY708 | | | | | | | | | 127 |
| NY717 | | 0.8 | 15 | | 28 | -2.6 | 11.5 | -1.6 | 102 |
| NY723 | 48 | -1.11 | 8 | -5.73 | 34 | | 22.5 | | |
| NY733 | | | | | | | | | |
| NY736 | 44 | -1.7 | 15 | 0.6 | 36 | 0.7 | 11.5 | -3.2 | 90 |
| NY737 | | -0.7 | 14 | -0.3 | 33 | -1.5 | 19.5 | -1.6 | 90 |
| NY741 | 0 | -7.7 | 14 | -0.3 | 24.5 | -7.3 | 11 | -3.8 | 99 |
| NY743 | 53 | -0.4 | 15 | 0.6 | 35 | | | | 97 |
| NY748 | 55 | 0.2 | 15 | 0.7 | 35 | -0.1 | 21 | -0.9 | 91 |

**Table S1. Subject Clinical Neuropsychological Characteristics.** Raw and normalized results for Rey Auditory Verbal Learning Test (RAVLT), a standard neuropsychological assessment of verbal memory are provided for 15 subjects. Raw and normalized results for Rey-Osterreith Copy Figure Test (ROCFT), a standard neuropsychological assessment of non-verbal or visual memory is provided. Finally, test of premorbid functioning (TOPF) is a pre-injury measure of IQ score. Missing values means that neuropsychological testing data was not collected for that subject. Other subject demographic and clinical characteristics are reported in Table 1.

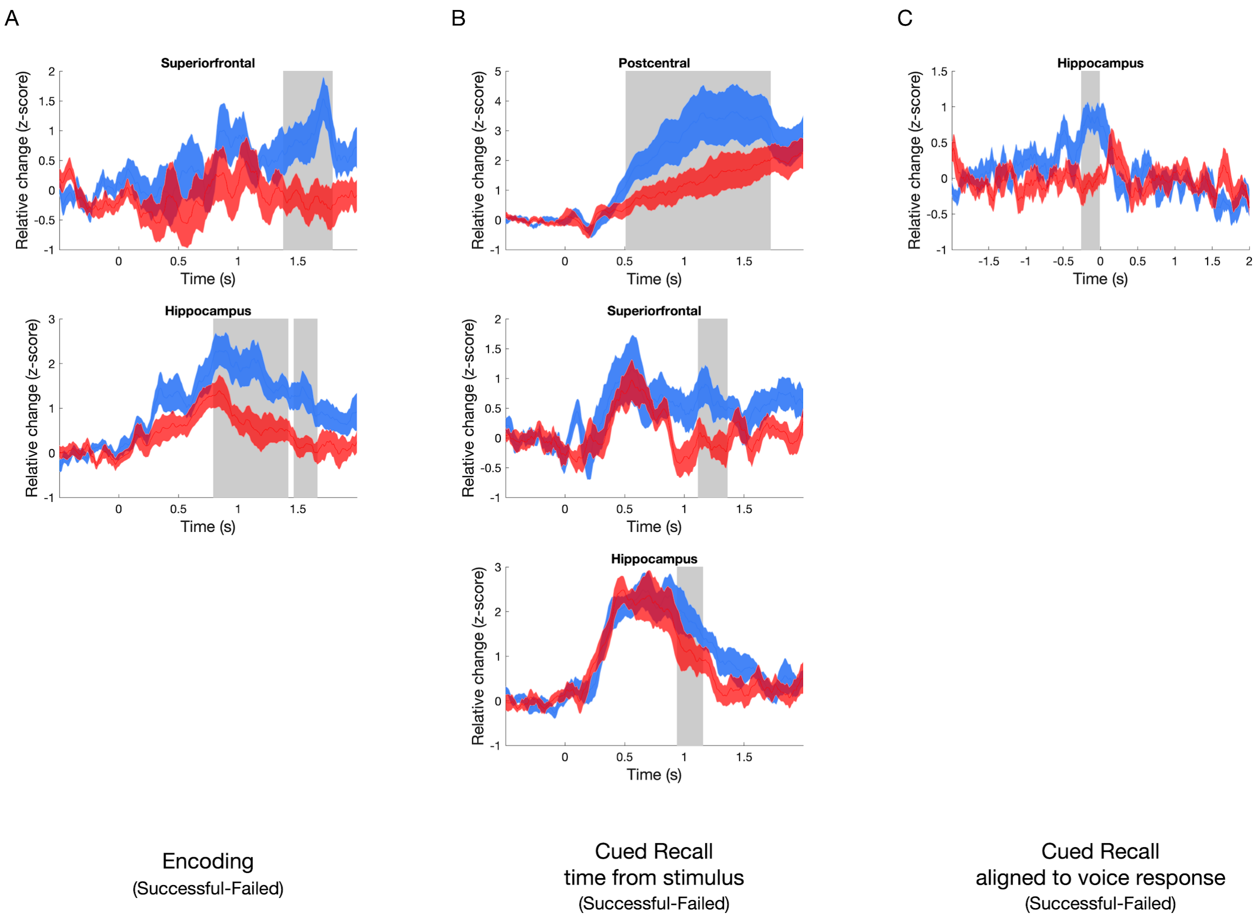


**Figure S1. Brain regions showing significant differences in the time-course of high-gamma activity (HGA) between successful and failed trials during** (A) Encoding, (B) Recall (C) Recall timed to response vocalization. All significant clusters identified at a significance threshold $p<0.05$ using a cluster-based permutation test.

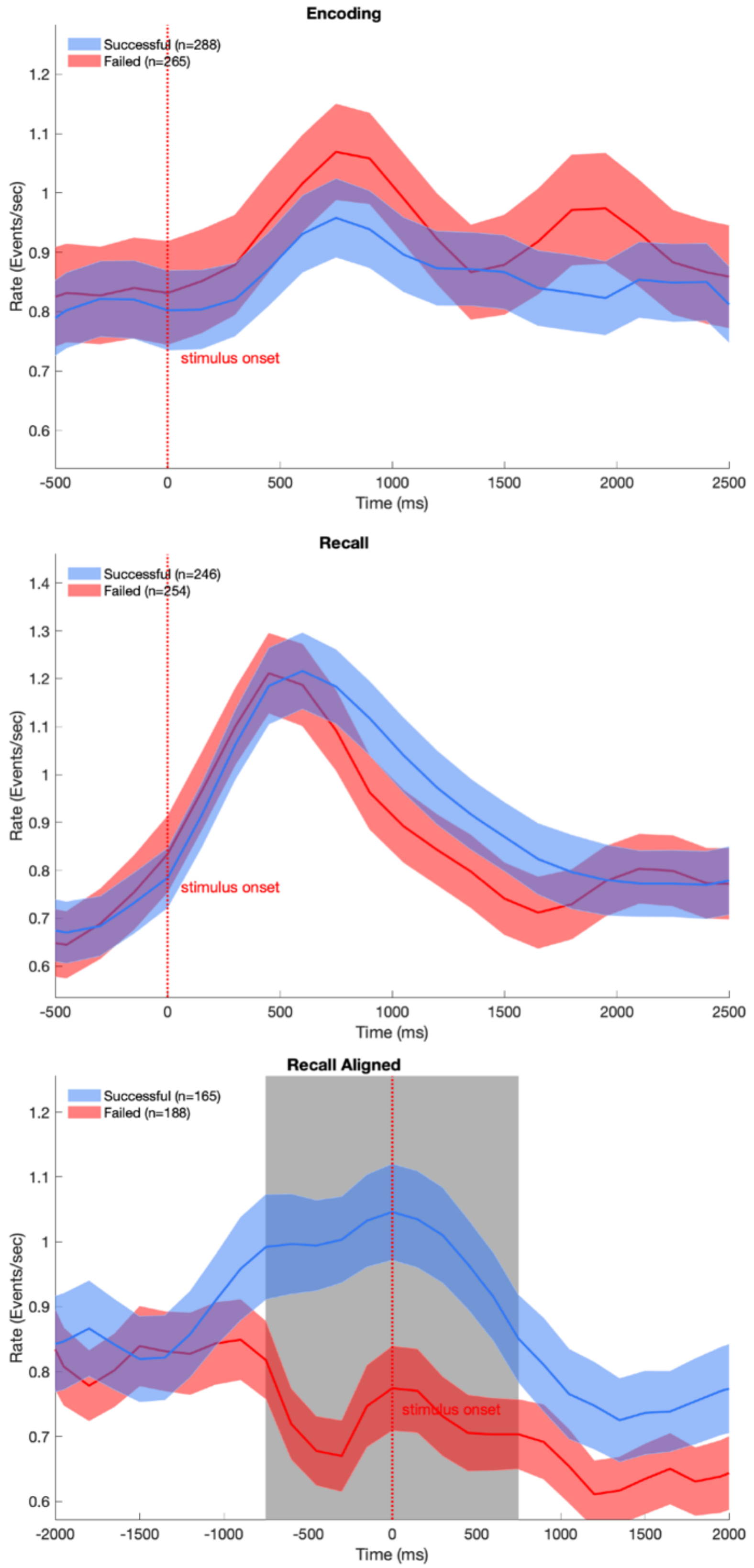


**Figure S2. Difference in Hippocampal Ripple Rate between Successful and Failed Associative Memory Trials, within the seizure onset zone (SOZ).** In the primary analysis, only hippocampal electrodes outside of the SOZ were included. Seven patients had hippocampal electrodes within the SOZ, allowing comparison of physiological response during the memory task. **TOP.** No difference in hippocampal ripple rate for electrodes within SOZ between successful and failed encoding trials. MIDDLE. No difference in hippocampal ripple rate for electrodes within SOZ between successful and failed cued recall trials. BOTTOM. When aligned to the voice response, a high hippocampal ripple rate distinguishing between successful and failed trials, occurring from -800 ms to +700 ms (BLUE, $p<0.05$, cluster-corrected) to voice response, compared to failed trials (RED). These results suggest diminished neurophysiological response during memory processes for hippocampal brain tissue within the SOZ.

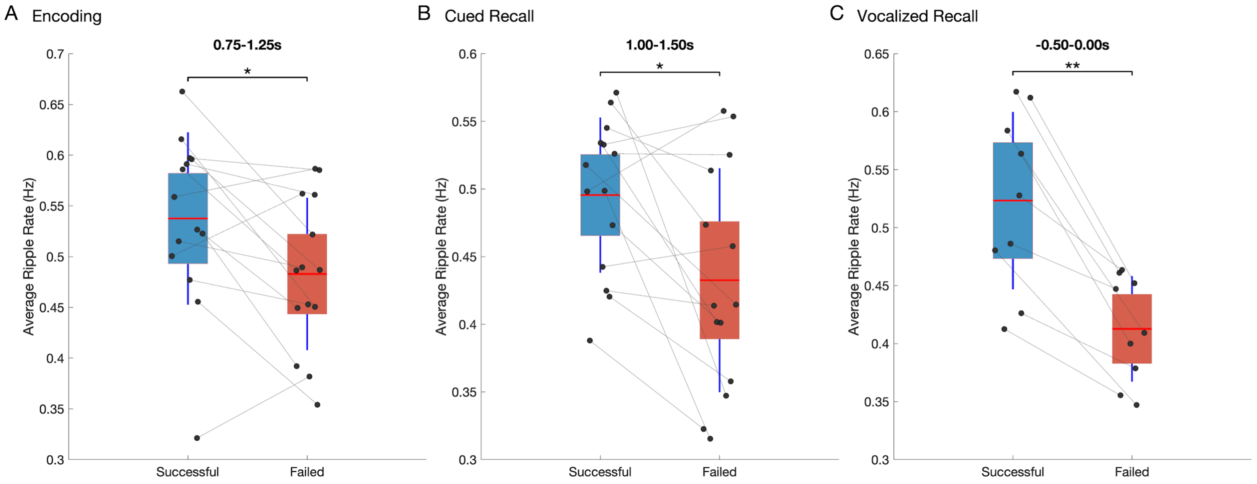


**Figure S3. Ripple rate differences between successful and failed associated memory trials.** Within-subjects difference in ripple rate for successful and failed trials during critical time windows for (A)encoding (0.75-1.25 sec, Z=-2.23, p=0.026, Wilcoxon signed rank test), (B) recall (1-1.5 sec, Z=-2.48, p=0.013), (C) vocalization-aligned recall (-0.5-0 sec, Z=-2.67, p=0.008). Ripple rate for successful and failed trials for each subject is connected by lines and shows that for most patients, ripple rate decreases during failed trials. Red line indicates group mean for each condition, red and blue boxes are the 95% confidence interval for successful and failed trials respectively and error bars present the standard deviation.

A. Encoding

B. Recall

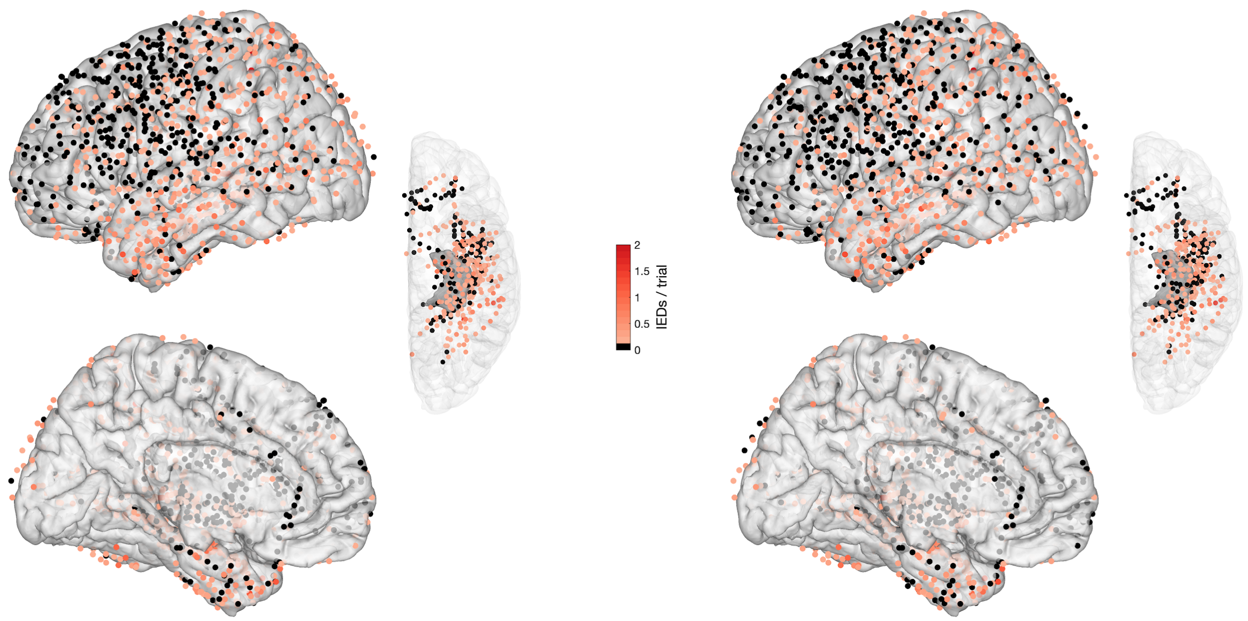


**FIgure S5. Distribution of IEDs across iEEG electrodes.** IED rate aggregated across all 15 Patients during (A) Encoding and (B) Recall. Electrodes are color coded by the average number of IEDs per trial. Electrodes without any recorded IEDs are shown in black.

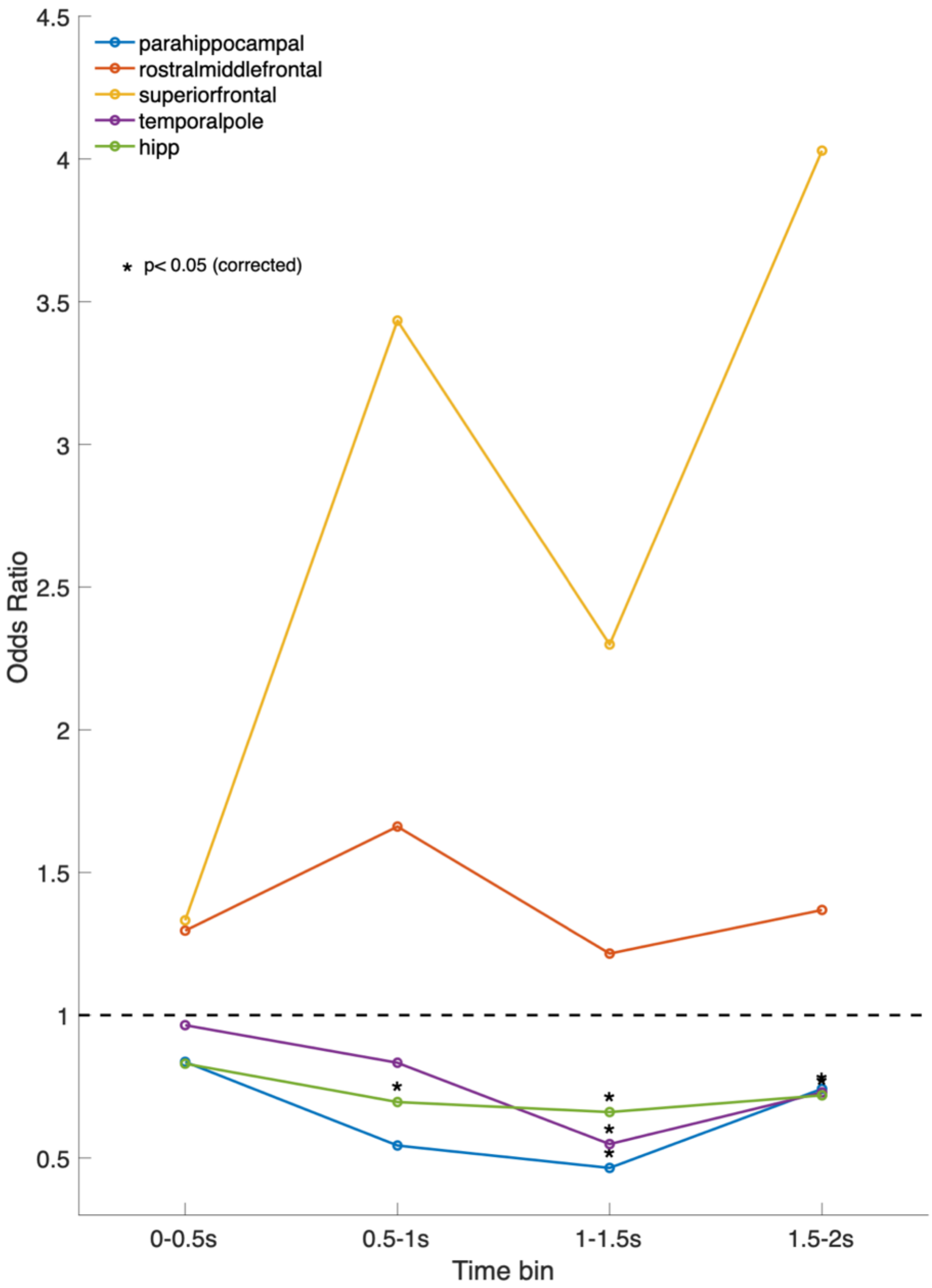


**FIgure S5. Odds of successful memory for IEDs occurring in selected frontal and temporal brain regions.** IED impact on odds of remembering was calculated for brain regions which demonstrated a critical difference in HGA (From **Figure 2)**. IEDs in frontal regions did not have a negative impact on memory performance. On the other hand, IEDs in medial temporal lobe and temporal pole were associated with decreased odds of remembering. After correction for multiple comparisons, significant changes are indicated by an asterisk located in hippocampus, temporal pole, and parahippocampal regions (FDR correction p<0.05).

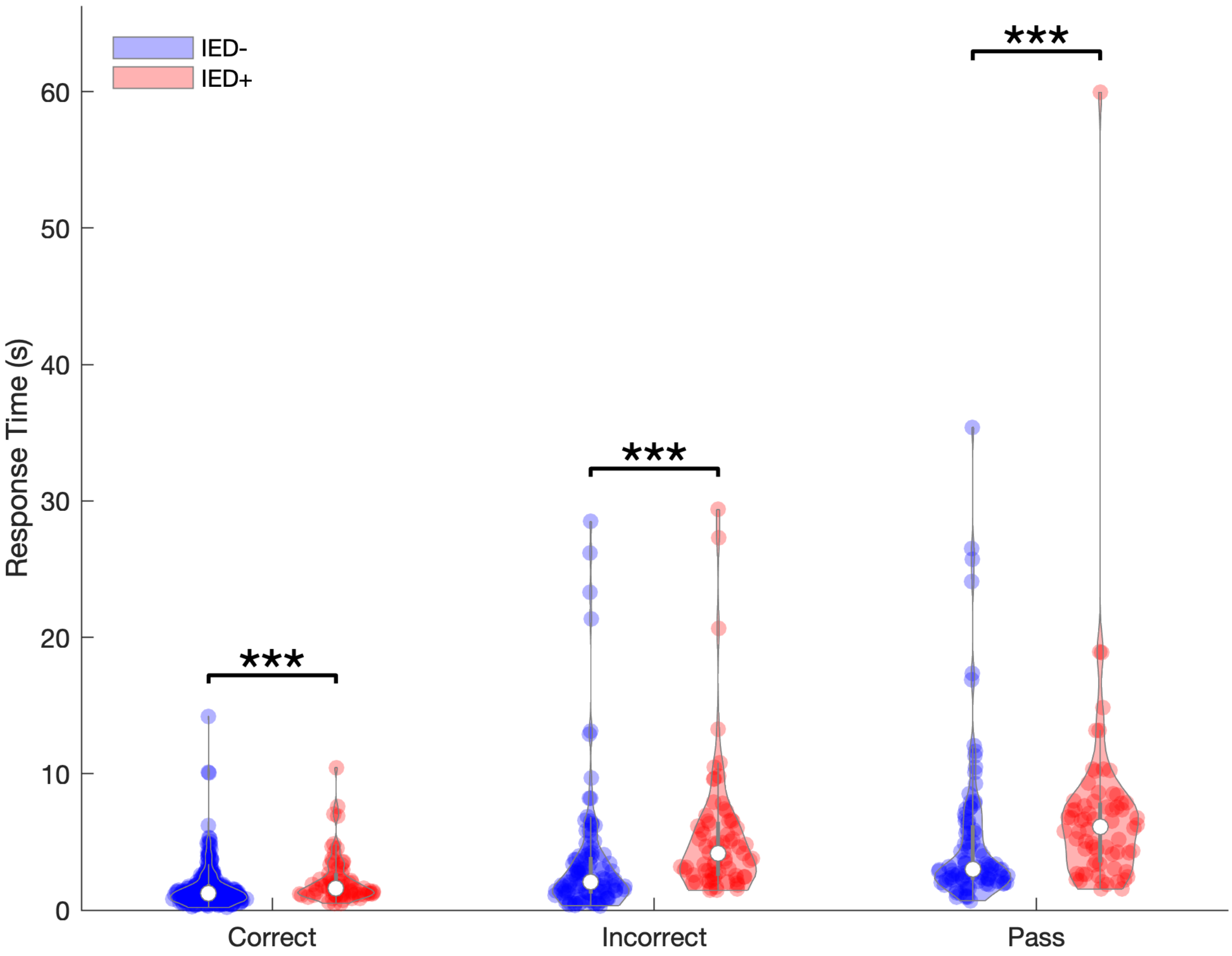


**FIgure S6. Trials with IEDs demonstrated slower response times than trials without IEDs**. Trials with IEDs (RED) were significantly slower than for trials without an IED (BLUE), for correctly vocalized responses, incorrectly vocalized responses, and passes. Response times were calculated from time of cue presentation to onset of vocalization, with the latter manually scored. Comparison of response latencies between trials with an IED (RED) and without an IED (BLUE) were conducted using Wilcoxon's rank-sum test (*** $p<0.001$). For all response types (correct, incorrect, pass), trials with an IED were slower compared to trials without an IED when trials were pooled across patients (**Correct:** $median_{IED-}$ = 1.25 (1.10 IQR); $median_{IED+}$ = 1.61 (1.49 IQR), Z = -3.89, $p<0.001$; **Incorrect:** $median_{IED-}$ = 2.05 (2.62 IQR); $median_{IED+}$ = 4.15 (3.93 IQR), Z = -5.19, $p<0.001$, **Pass:** $median_{IED-}$ = 3.01 (4.03 IQR); $median_{IED+}$ = 6.13 (4.35 IQR), Z=-3.74, $p<0.001$; Wilcoxon rank sum test).

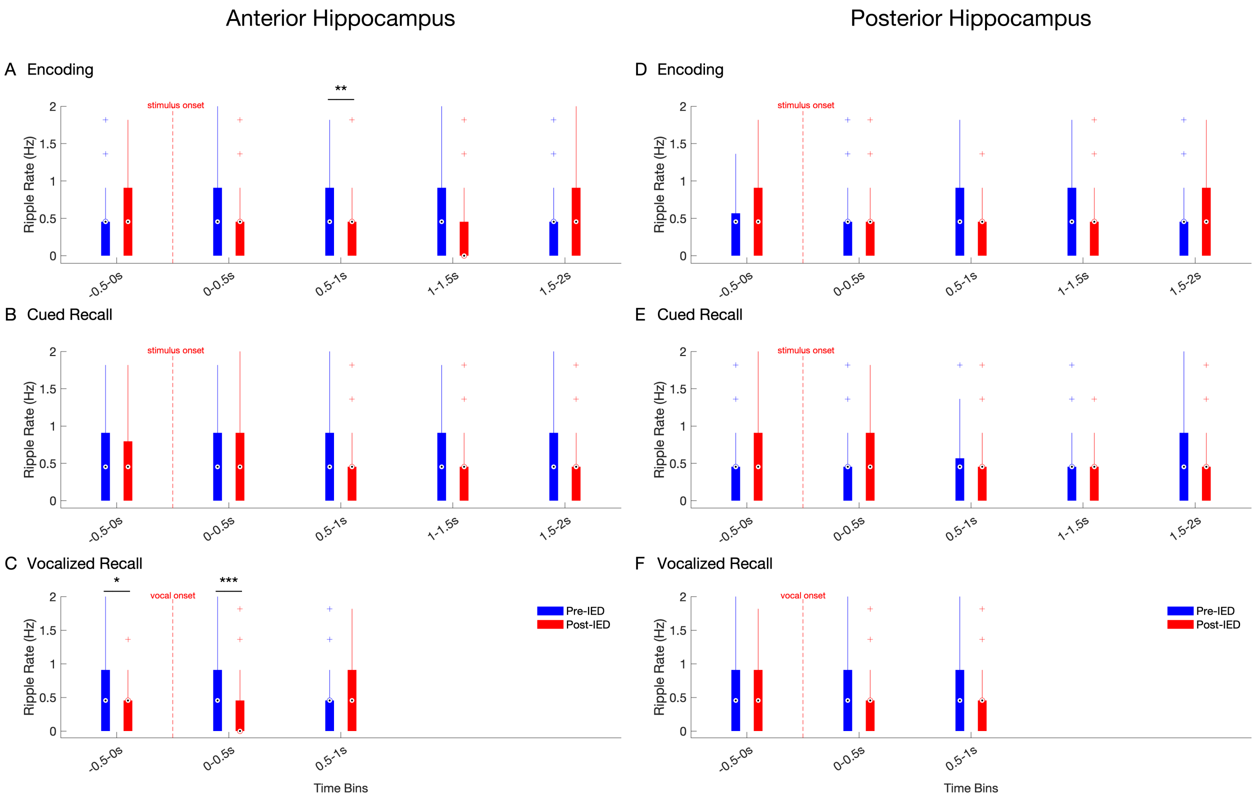


**Fig. S7 Average Ripple Rate during Encoding & Recall in Anterior and Posterior Hippocampus.** Ripple rate is calculated in 500ms pre- (blue) vs. 500ms post-IED (red) bins, time-locked to IEDs detected in 500 ms bins during Encoding, Cued Recall, and Voice-aligned Recall. Box and whisker plots represent the median (circles) and interquartile range (bars), along with extreme values (whiskers and outliers). Findings are calculated separately for Anterior Hippocampus (A-C) and Posterior Hippocampus (D-F). A reduction in ripple rate after an IED event was found during the 0.5-1s ($Z=2.2921$, $N_{IEDs}=119$, $p=0.02$, Wilcoxon signed-rank test) a encoding window only in anterior hippocampus. In addition, ripple rates were significantly reduced in the time windows immediately preceding (-0.5-0s, $Z=2.0818$, $N_{IEDs}=103$, $p=0.03$) and during (0-0.5s, $Z=3.1518$, $N_{IEDs}=86$, $p=0.002$) vocalization during cued recall only in anterior hippocampus.
